# A toolbox for imaging RIPK1, RIPK3 and MLKL in mouse and human cells

**DOI:** 10.1101/2020.10.26.356063

**Authors:** André L. Samson, Cheree Fitzgibbon, Komal M. Patel, Joanne M. Hildebrand, Lachlan W. Whitehead, Joel S. Rimes, Annette V. Jacobsen, Christopher R. Horne, Xavier J. Gavin, Samuel N. Young, Kelly L. Rogers, Edwin D. Hawkins, James M. Murphy

## Abstract

Necroptosis is a lytic, inflammatory cell death pathway that is dysregulated in many human pathologies. The pathway is executed by a core machinery comprising the RIPK1 and RIPK3 kinases, which assemble into necrosomes in the cytoplasm, and the terminal effector pseudokinase, MLKL. RIPK3-mediated phosphorylation of MLKL induces oligomerization and translocation to the plasma membrane where MLKL accumulates as hotspots and perturbs the lipid bilayer to cause death. The precise choreography of events in the pathway, where they occur within cells, and pathway differences between species, are of immense interest. However, they have been poorly characterized due to a dearth of validated antibodies for microscopy studies. Here, we describe a toolbox of antibodies for immunofluorescent detection of the core necroptosis effectors, RIPK1, RIPK3 and MLKL, and their phosphorylated forms, in human and mouse cells. By comparing reactivity with endogenous proteins in wild-type cells and knockout controls in basal and necroptosis-inducing conditions, we characterise the specificity of frequently-used commercial and recently-developed antibodies for detection of necroptosis signaling events. Importantly, our findings demonstrate that not all frequently-used antibodies are suitable for monitoring necroptosis by immunofluorescence microscopy, and methanol-is preferable to paraformaldehyde-fixation for robust detection of specific RIPK1, RIPK3 and MLKL signals.

## INTRODUCTION

Cell death by necroptosis is thought to have originated as an ancestral host defence mechanism, which is reflected in the breadth of pathogen-encoded proteins that inhibit the pathway^1, 2, 3, 4, 5, 6^. In addition to reported innate immunity functions^7, 8, 9^, the dysregulation of necroptosis has been implicated in a range of pathologies, including ischemic-reperfusion injuries, such as in the kidney^10, 11^ and heart^12^, and inflammatory diseases^13, 14, 15, 16^, including inflammatory bowel disease^17^. Accordingly, there is widespread interest in therapeutically-targeting the pathway to counter human disease. Owing to the recent identification of the terminal effectors of the pathway, RIPK3 (in 2009)^1, 18, 19^ and MLKL (in 2012)^20, 21^, however, the extent of indications attributable to necroptotic cell death is poorly understood. Precisely defining pathologies impacted by necroptosis has posed a challenge owing to the dearth of antibodies validated to specifically detect members of the pathway and their activated (phosphorylated) forms in fixed cells and tissues. As a result, the contribution of necroptosis to many pathologies remains a subject of ongoing debate^22, 23, 24, 25^.

Necroptotic cell death signaling is initiated by ligation of death receptors, such as the TNF receptor 1, or pathogen detectors, such as Toll-like receptors 3 and 4 and the ZBP1/DAI intracellular viral RNA sensor protein. In cellular contexts where the activities of the Inhibitors of Apoptosis proteins (IAPs) E3 Ubiquitin ligase family and the proteolytic apoptotic effector, Caspase-8, are depleted or compromised, necroptosis ensues. The precise choreography of necroptotic signaling is still emerging, although recent studies have defined key events and checkpoints in the pathway^26, 27, 28, 29^. Following pathway induction, RIPK1 autophosphorylation prompts hetero-oligomerization with RIPK3 via an amyloid-forming motif in the region C-terminal to their kinase domains termed the RHIM (RIP Homotypic Interaction Motif) to form a cytoplasmic platform known as the necrosome^30,31,32^. Upon RIPK3 activation by autophosphorylation within necrosomes^33, 34^, RIPK3 is primed to phosphorylate the activation loop of the MLKL pseudokinase domain to activate MLKL’s killing function^14, 20, 27, 28, 35, 36, 37, 38, 39^. In the case of human MLKL, dormant MLKL appears to be at least in part pre-associated with RIPK3 in the cytoplasm^20, 40^, and stable recruitment to necrosomes appears to be an essential checkpoint in MLKL activation^28^. In the case of mouse MLKL, a transient interaction between RIPK3 and MLKL appears to be sufficient for MLKL activation in mouse cells^35, 36, 37^. Regardless of species, MLKL phosphorylation is thought to provoke a conformational change in the pseudokinase domain that leads to oligomerization and unmasking of the killer N-terminal four-helix bundle (4HB) domain^37, 40, 41, 42, 43^. The human MLKL 4HB domain likely engages chaperones to facilitate translocation to the plasma membrane via an actin-, Golgi- and microtubule-dependent mechanism, where MLKL accumulates in hotpots. When a threshold is surpassed, the 4HB domains of MLKL permeabilize the membrane to induce cell death^26, 29^.

While biochemical studies have defined these steps and checkpoints, visualizing the spatiotemporal dynamics of endogenous proteins during in necroptosis using microscopy-based approaches has proven challenging in the absence of antibodies that have been validated for target specificity. Similarly, the lack of validated reagents poses a challenge to immunohistochemical staining of patient tissue sections, and therefore attribution of a role for necroptosis in pathologies, because knockout tissue controls are not available. Here, we have established procedures for staining endogenous RIPK1, RIPK3 and MLKL, and their phosphorylated forms, in fixed mouse and human cells. While several frequently-used antibodies were found to be suitable for selectively staining these proteins, as validated by comparisons with cells deficient for each protein, many exhibited non-specific staining and are therefore unsuitable for immunofluorescence and immunohistochemistry. Our studies also highlight the importance of validating antibody compatibility with fixation methods. In most cases, paraformaldehyde fixation ablated epitopes, presumably because RIPK3 and MLKL are lysine-rich and prone to modification by crosslinking, whereas methanol fixation enabled specific detection of these proteins. Due to the sequence divergence between mouse and human RIPK3 and MLKL^41, 43, 44, 45, 46^, it was not possible to specifically-detect proteins across species using a single reagent, which necessitated the development of a new antibody that specifically detected mouse MLKL. Collectively, our studies present a toolbox of selective antibodies that will enable critical analysis of the chronology, checkpoints and kinetics of necroptotic signaling in mouse and human cells.

## RESULTS

### Immunofluorescent detection of human MLKL

We recently described a monoclonal antibody, clone 10C2 (source: WEHI Antibody Facility, in-house), that recognises an epitope centred on residues 413-471 of human MLKL^29^ (Fig. 1a). The initial screen that identified this clone suggested it yielded specific immunosignals when cells were fixed with methanol, but not when cells were fixed with paraformaldehyde. Organic solvents such as methanol preserve cells by precipitating proteins, whereas aldehyde-based agents such as paraformaldehyde fix cells by crosslinking lysines and other primary amines. Because human MLKL is lysine-rich (93^rd^ percentile in the cytoplasmic and membrane-associated proteome), paraformaldehyde likely causes widespread crosslinking of MLKL, which in turn masks epitopes and reduces immunoreactivity. As several other necroptotic proteins are lysine-rich, we considered whether the choice of fixative was a critical variable for robust immunodetection of MLKL, RIPK3 and RIPK1 in human and mouse cells. Accordingly, we compared the performance of 22 antibodies for immunofluorescent staining of human HT29 cells and mouse dermal fibroblasts (MDFs) - two cellular models that are well-characterized to undergo necroptosis when treated with TNF, Smac-mimetic and IDN-6556 (herein referred to as TSI)^5, 28, 29, 44, 47^. To quantitatively gauge the performance of each antibody, their immunosignals were characterized in four ways: 1) A ratio of the immunofluorescent signals between a positive and negative control. This produces a signal-to-noise curve that estimates ‘specific signal abundance’ (Fig. 1b). For testing phospho-specific antibodies, wild-type cells undergoing necroptosis were used as a positive control and untreated wild-type cells as a negative control (Fig. S1). For testing all other antibodies, untreated wild-type cells were used as a positive control and relevant knockout cells as a negative control. 2) Micrographs were gated so that the minimum/maximum immunosignals corresponded to the 5^th^/95^th^ percentiles of the signal-to-noise curve (Fig. 1b). 3) The percentage of total signals that fall within this gate were determined to show how much signal from the positive control was considered specific (Fig. 1c). 4) Antibodies were immunoblotted against positive and negative controls to independently assess their specificity.

**Figure 1:**
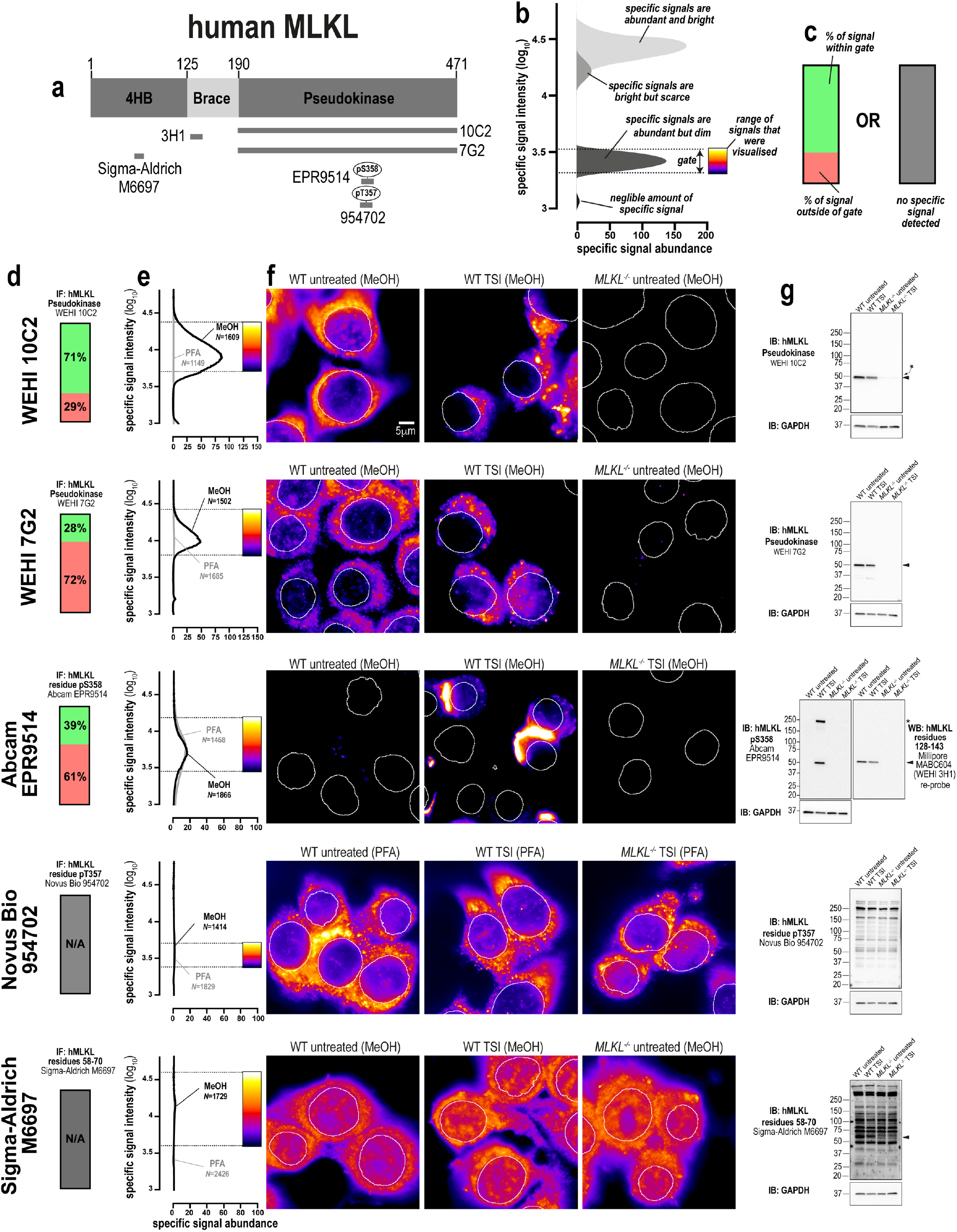
Methanol fixation is optimal for the immunofluorescent detection of human MLKL. **a** Human MLKL domain architecture showing the immunogens used to raise the tested anti-MLKL antibodies. **b** Demonstration of how signal-to-noise ratios were used to quantify the abundance and brightness of specific immunofluorescent signals generated by different antibodies. The 5^th^ and 95^th^ percentile of each signal-to-noise curve defines the gate where specific immunosignals were observed. As indicated by the pseudocolour look-up-table, only immunosignals within this gate were visualised. **c** Chart exemplifying how the amount of signal within the gate, relative to the total amount of detectable signal, provides another gauge of antibody specificity for immunofluorescence. **d** Quantitation of the percentage of gated signals for the tested MLKL antibodies. **e** Quantitation of specific signal abundance produced by the tested MLKL antibodies on methanol-fixed (MeOH) or paraformaldehyde-fixed (PFA) HT29 cells. The number of cells imaged (N) to generate each signal-to-noise curve is shown. **f** Micrographs of immunofluorescent signals for the tested MLKL antibodies on HT29 cells. As indicated by each pseudocolour look-up-table, only immunosignals within the respective gate in panel **e** were visualised. Data are representative of n=3 (clones 10C2, 7G2) and n=2 (Abcam clone EPR9514, Novus Biological MAB9187/clone 954702 and Sigma-Aldrich M6697) independent experiments. Nuclei were detected by Hoechst 33342 staining and are demarked by white outlines in micrographs. **g** Immunoblot using the tested MLKL antibodies against wild-type and *MLKL*^*-/-*^ HT29 cell lysates. Closed arrowheads indicate the main specific band. Asterisks indicate non-specific bands that could otherwise confound data interpretation. Immunoblots were re-probed for GAPDH as loading control.

Using this approach, we confirmed that clone 10C2 specifically detects endogenous human MLKL in methanol-fixed, but not paraformaldehyde-fixed cells (Fig. 1d-f). The 10C2 clone produces a favourable immunostaining profile, where the specific signals are abundant (Fig. 1e) and these specific signals represent the majority of all detectable signals (Fig.1d). We recently developed another MLKL-specific antibody, clone 7G2 (source: in-house), that recognises an epitope centred on *α*F-*α*G loop the C-lobe of the human MLKL pseudokinase domain, which differs to the site recognised by the 10C2 clone^29^. Despite binding a distinct site in human MLKL to 10C2, the 7G2 clone shows similar specificity for human MLKL in immunoblot analyses and only yields specific immunofluorescent signals on methanol-fixed cells (Fig. 1d-f). Notably, clone 7G2 is inferior to clone 10C2 for immunofluorescence studies, because specific signals were less abundant (Fig. 1e) and were a minor fraction of all detectable signals (Fig.1d).

We observed that clone EPR9514 (source: Abcam), an antibody raised against phospho-S358 of human MLKL^48^, which is a hallmark of MLKL activation during necroptosis^20^, produced specific and comparable immunofluorescent signals in both methanol- and paraformaldehyde-fixed cells undergoing necroptosis (Fig. 1e). As MLKL can be disulfide-crosslinked during necroptosis^49, 50^ (Fig. S2A-B), we tested whether fixation in the presence of *N*-ethylmaleimide (NEM) to prevent further disulfide bonding, or post-fixative treatment with Tris(2-carboxyethyl)phosphine (TCEP) to reduce disulfide bonds, altered the immunofluorescent detection of phospho-MLKL by clone EPR9514. However, neither preventing nor reducing disulfide bonds in fixed cells precluded the immunodetection of MLKL by clone EPR9514 (Fig. S2C-D). Indeed, disulfide-crosslinking may be a by-product of MLKL activation given that recombinant human MLKL oligomerizes in the absence of disulfide bonds (Fig. S2E).

Two other anti-human MLKL antibodies were found to be non-specific (Novus Biological clone 954702; and Sigma-Aldrich M6697, a polyclonal antibody raised against residues 58-70; Fig. 1a and 1e), with equivalent signal intensity and diffuse staining observed in wild-type and *MLKL*^*-/-*^ cells. Lastly, immunoblotting confirmed that clones 10C2, 7G2 (both in-house) and EPR9514 (Abcam) were highly-specific with bands corresponding to human MLKL’s molecular weight of 54kDa observed in wild-type but not *MLKL*^*-/-*^ HT29 cell lysates. In contrast, clone 954702 (Novus Biological MAB9187) and the Sigma-Aldrich M6697 polyclonal antibody were non-specific, with multiple bands not corresponding to the molecular weight of MLKL observed in both wild-type and *MLKL*^*-/-*^ HT29 cell lysates (Fig. 1g).

In summary, three of the five antibodies that were tested selectively recognised human MLKL in methanol-fixed cells (clones 10C2, 7G2 and EPR9514) and, accordingly, we recommend methanol fixation for detecting endogenous human MLKL by immunofluorescence. Under these conditions, cytoplasmic forms of MLKL could be detected using the 10C2 and 7G2 clones, and phospho-MLKL accumulating at the cell periphery with the EPR9514 clone (Fig. 1f). These data underscore the importance of careful consideration of fixation method when using MLKL as a histological marker of necroptosis.

### Immunofluorescent detection of human RIPK3

We next assessed five anti-human RIPK3 antibodies (Fig. 2a). Two antibodies raised against the C-terminus (source: ProSci 2283, Novus Biological NBP2-24588) and one antibody raised against the N-terminus of RIPK3 (source: Novus Biological MAB7604, clone 780115) yielded non-specific signals via both immunofluorescence and immunoblotting (Fig. 2b-e). Our recently described antibody, clone 1H2 (source: in-house)^5^, detected RIPK3 via immunofluorescence in methanol-fixed, but not paraformaldehyde-fixed cells. However, most of the immunofluorescent signals produced by 1H2 were non-specific (Fig. 2b), and the remaining signals are specific but of very low abundance (Fig. 2c). Thus, although clone 1H2 is highly-selective when used as an immunoblotting reagent (Fig. 2d^5^), it should only be used for the immunofluorescent detection of human RIPK3 under carefully controlled conditions. Our data also showed that clone D6W2T (source: Cell Signaling Technology), an antibody raised against a mark of human RIPK3 activation in necroptosis, phospho-S227, produced specific immunofluorescent signals in paraformaldehyde-fixed cells undergoing necroptosis, but not in methanol-fixed cells (Fig. 2d). Again, the low abundance of these specific immunosignals suggests that clone D6W2T should only be used for immunofluorescence under carefully controlled conditions, such as where *RIPK3*^*-/-*^ control cells are available for direct comparison. Another caveat is that immunoblotting showed that clone D6W2T not only detected RIPK3 in cells undergoing necroptosis, but also in untreated cells when RIPK3 was presumably not phosphorylated at S227 (Fig. 2d). Therefore, while two of the five antibodies that were tested specifically recognised human RIPK3, the low abundance of their specific signals complicates use of these antibodies for studying endogenous human RIPK3 via immunofluorescence. At present, more specific antibodies are needed to enable the immunofluorescent detection of human RIPK3.

**Figure 2:**
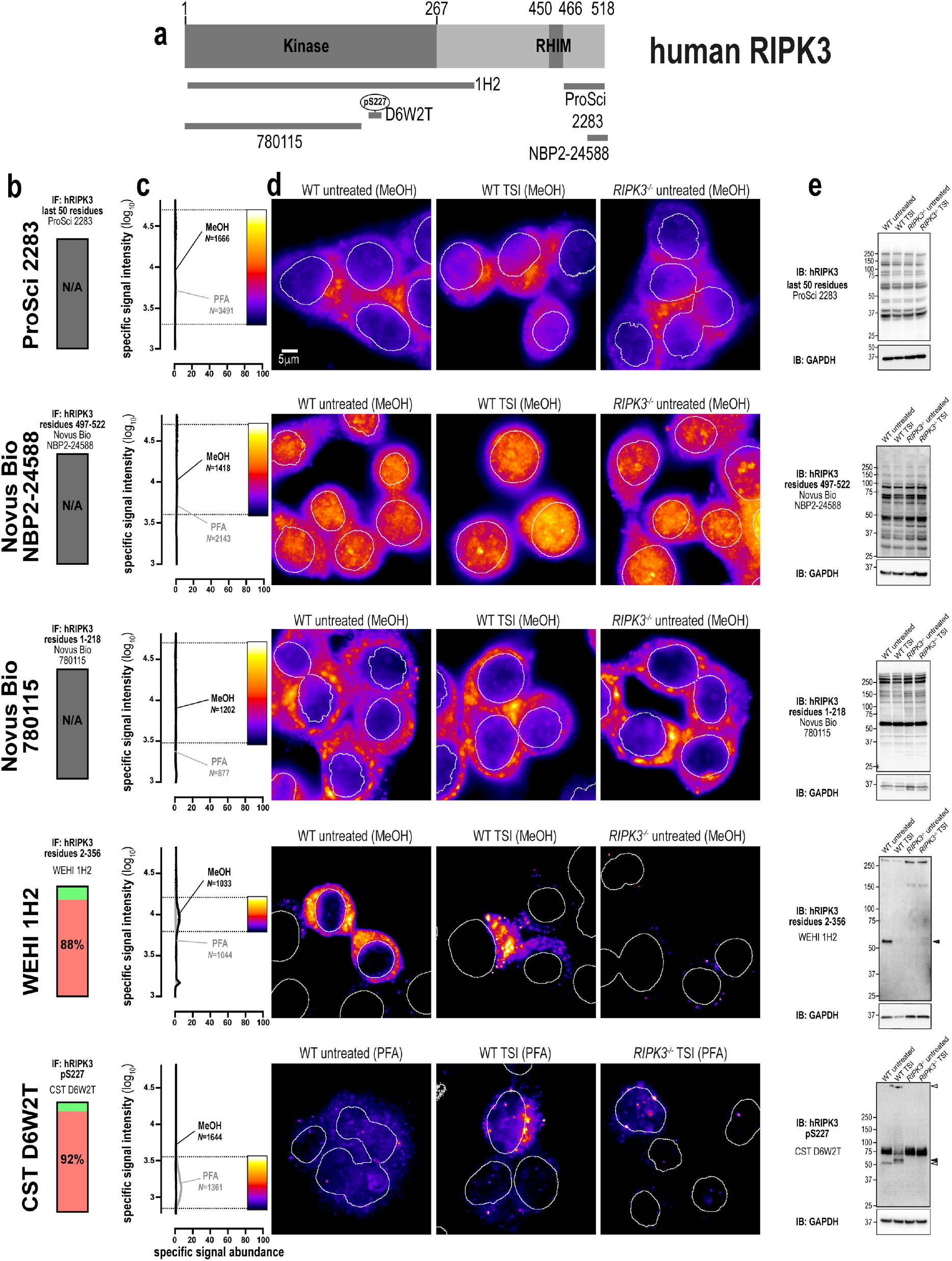
Better antibodies are needed for imaging human RIPK3. **a** Human RIPK3 domain architecture showing the immunogens or epitopes for the tested anti-RIPK3 antibodies. **b** Quantitation of the percentage of gated signals for the tested RIPK3 antibodies. **c** Quantitation of specific signal abundance produced by the tested RIPK3 antibodies on methanol-fixed (MeOH) or paraformaldehyde-fixed (PFA) HT29 cells. The number of cells imaged (N) to generate each signal-to-noise curve is shown. **d** Micrographs of immunofluorescent signals for the tested RIPK3 antibodies on HT29 cells. As indicated by each pseudocolour look-up-table, only immunosignals within the respective gate in panel **c** were visualised. Data are representative of n=3 (Cell Signaling Technology clone D6W2T and in-house clone 1H2) and n=2 (ProSci 2283, Novus Biological NBP2-24588 and MAB7604/clone 780115) independent experiments. Nuclei were detected by Hoechst 33342 staining and are demarked by white outlines in micrographs. **e** Immunoblot using the tested RIPK3 antibodies against wild-type and *RIPK3*^*-/-*^ HT29 cell lysates. Closed arrowheads indicate the main specific band. Open arrowheads indicate other specific bands of interest. Immunoblots were re-probed for GAPDH as loading control.

### Immunofluorescent detection of human RIPK1

We next tested three monoclonal anti-RIPK1 antibodies: one against the N-terminal domain (source: Cell Signaling Technology clone D94C12), one against the C-terminal region (source: BD Transduction Laboratories clone 38/RIP) and one raised against phospho-S166 (source: Cell Signaling Technology clone D8I3A; Fig. 3a). All three clones yielded specific immunofluorescent signals in methanol- and paraformaldehyde-fixed cells, however, these signals were more abundant in methanol-fixed samples (Fig. 3b-d). All three clones were also specific via immunoblot (Fig. 3e).

**Figure 3:**
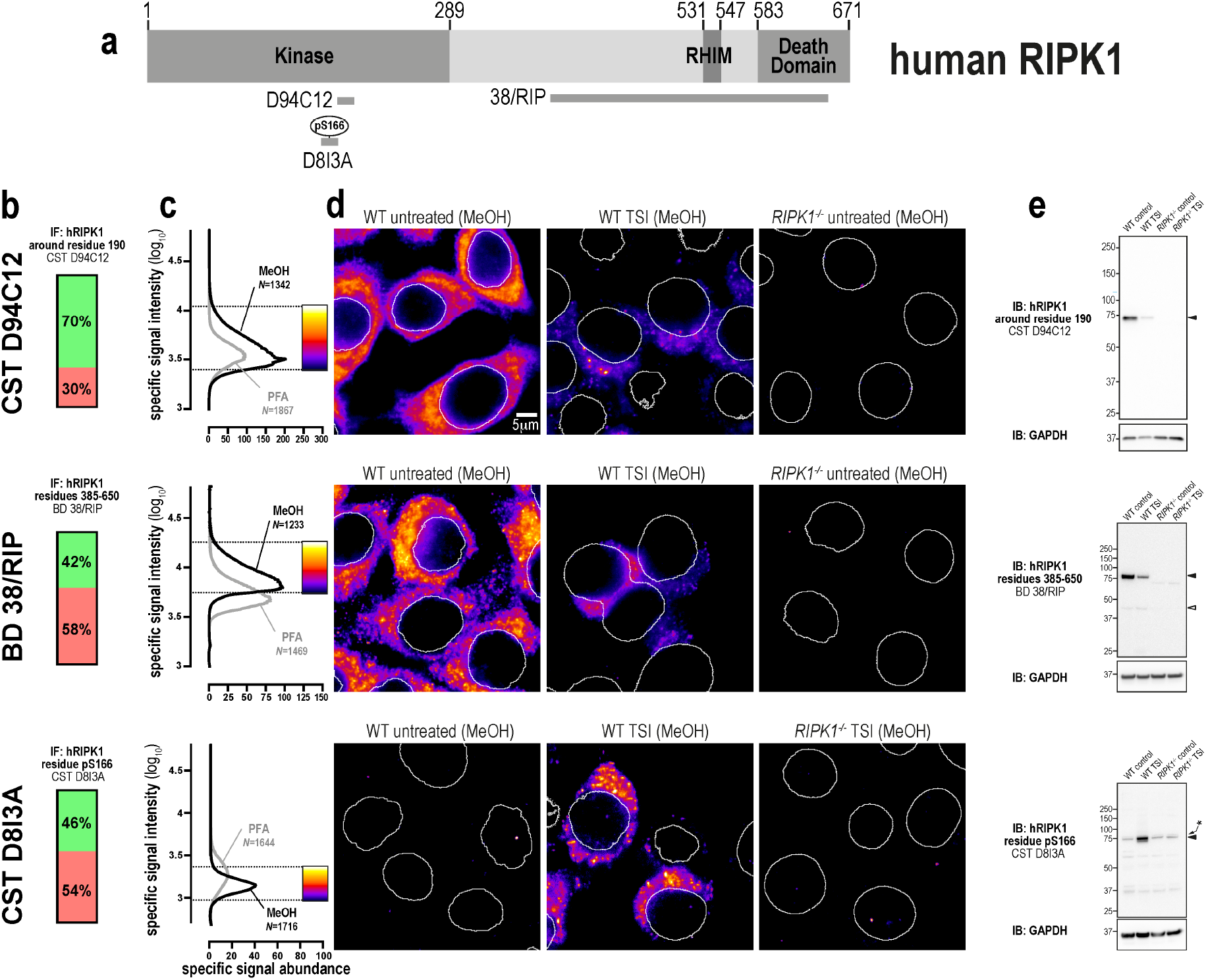
Three specific antibodies for imaging endogenous human RIPK1. **a** Human RIPK1 domain architecture showing the immunogens or epitopes for the tested anti-RIPK1 antibodies. **b** Quantitation of the percentage of gated signals for the tested RIPK1 antibodies. **c** Quantitation of specific signal abundance produced by the tested RIPK1 antibodies on methanol-fixed (MeOH) or paraformaldehyde-fixed (PFA) HT29 cells. The number of cells imaged (N) to generate each signal-to-noise curve is shown. **d** Micrographs of immunofluorescent signals for the tested RIPK1 antibodies on HT29 cells. As indicated by each pseudocolour look-up-table, only immunosignals within the respective gate in panel **c** were visualised. Data are representative of n=2 (Cell Signaling Technology clones D94C12 and D8I3A, BD Transduction laboratories clone 38/RIP) independent experiments. Nuclei were detected by Hoechst 33342 staining and are demarked by white outlines in micrographs. **e** Immunoblot using the tested RIPK1 antibodies against wild-type and *RIPK1*^*-/-*^ HT29 cell lysates. Closed arrowheads indicate the main specific band. Open arrowheads indicate other specific bands of interest. Asterisks indicate non-specific bands that could otherwise confound data interpretation. Immunoblots were re-probed for GAPDH as loading control.

Importantly, the immunosignals from clones D94C12 and 38/RIP were markedly lower in cells undergoing necroptosis than in unstimulated wild-type cells (Fig. 3d-e). This observation is likely due to RIPK1 undergoing widespread post-translational modification and/or proteasomal degradation during TNF-induced cell death^51, 52^. Thus, despite clones D94C12 and 38/RIP exhibiting a favourable immunostaining profile in unstimulated cells, their decreased reactivity in cells undergoing necroptosis indicates that they should be used judiciously for immunofluorescent labelling of samples for accurate analysis of necroptosis. Conversely, while clone D813A preferentially stains RIPK1 in cells undergoing necroptosis, specific signals that were detected also had low intensity (Fig. 3b) and low abundance (Fig. 3c). In summary, three highly-specific antibodies exist for human RIPK1. However, because their specific signals have very low intensity and abundance in necroptotic cells, their use for the immunofluorescent detection of RIPK1 during necroptosis should be carefully controlled, including side-by-side examination of wild-type and *RIPK1*^*-/-*^ control cells.

### Immunofluorescent detection of mouse MLKL

To our knowledge, there are currently no validated monoclonal antibodies for the immunofluorescent detection for mouse MLKL in unstimulated cells. Accordingly, we raised a monoclonal antibody, clone 5A6 (source: in-house), against full-length mouse MLKL (Fig. 4a). The epitope for clone 5A6 resides in the C-terminal domain of mouse MLKL (Fig. S3). As shown in Fig. 4a-d, clone 5A6 produced specific and abundant signals for endogenous mouse MLKL in methanol-fixed, but not paraformaldehyde-fixed cells. Immunoblotting confirmed that clone 5A6 was highly-specific (Fig. 4e). Notably, compared to human MLKL which translocates en masse into cytoplasmic clusters during necroptosis, the relocation of mouse MLKL into cytoplasmic clusters during necroptosis was more subtle. This observation may relate to the notion that human MLKL is recruited to the necrosome in a more stable manner than mouse MLKL^28^.

**Figure 4:**
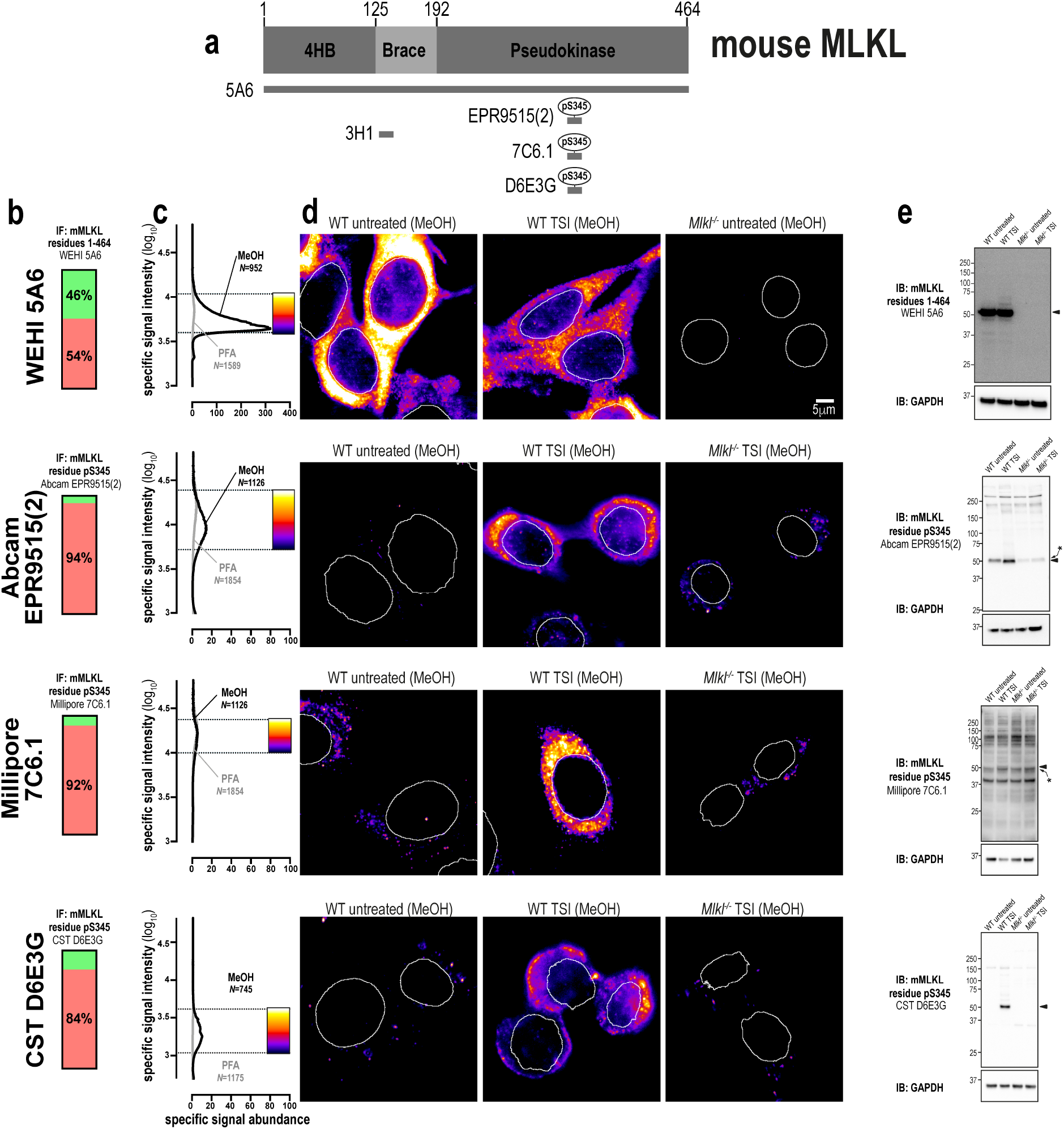
A new monoclonal antibody to image endogenous mouse MLKL. **a** Mouse MLKL domain architecture showing the immunogens or epitopes for the tested anti-MLKL antibodies. **b** Quantitation of the percentage of gated signals for the tested MLKL antibodies. **c** Quantitation of specific signal abundance produced by the tested MLKL antibodies on methanol-fixed (MeOH) or paraformaldehyde-fixed (PFA) MDFs. The number of cells imaged (N) to generate each signal-to-noise curve is shown. **d** Micrographs of immunofluorescent signals for the tested MLKL antibodies on MDFs. As indicated by each pseudocolour look-up-table, only immunosignals within the respective gate in panel **c** were visualised. Data are representative of n=3 (in-house clone 5A6) and n=2 (Abcam clone EPR9515(2), Millipore MABC1158/clone 7C6.1 and Cell Signaling Technology clone D6E3G) independent experiments. Nuclei were detected by Hoechst 33342 staining and are demarked by white outlines in micrographs. **e** Immunoblot using the tested MLKL antibodies against lysates from wild-type and *Mlkl*^*-/-*^ MDFs. Closed arrowheads indicate the main specific band. Asterisks indicate non-specific bands that could otherwise confound data interpretation. Immunoblots were re-probed for GAPDH as loading control.

Phosphorylation of mouse MLKL at S345 is essential for necroptosis in murine cells^37^, and therefore many antibodies have been raised against this phosphosite. We tested three commonly-used antibodies (Fig. 4a-e), namely clones: EPR9515(2) (source: Abcam), 7C6.1^39^ (source: Millipore MABC1158) and D6E3G (source: Cell Signaling Technology). All three clones yielded specific immunosignals in methanol-fixed cells (Fig. 4c), whereas only clone 7C6.1 was compatible with paraformaldehyde fixation. However, the performance of clone D6E3G was superior to the other anti-phospho-MLKL antibodies, because specific signals were more abundant (Fig. 4c) and represented a higher percentage of the total detectable signal (Fig. 4b). This trend was also evident via immunoblot, with clone 7C6.1 being largely non-specific, clone EPR9515(2) being moderately specific and clone D6E3G exhibiting a high degree of specificity towards activated MLKL in necroptotic cells (Fig. 4e).

In summary, clones 5A6 and D6E3G allow the immunofluorescent detection of mouse MLKL under resting and necroptotic conditions, respectively. Moreover, as with human MLKL, methanol is the fixative of choice for robust immunofluorescent detection of endogenous mouse MLKL.

### Immunofluorescent detection of mouse RIPK3

There are numerous well-validated antibodies raised against mouse RIPK3, including a polyclonal antibody against the C-terminus (source: ProSci 2283), and monoclonal antibodies recognising phospho-T231/phospho-S232 (source: Genentech clone GEN135-35-9^53^) and the extended kinase domain (residues 2-353) of mouse RIPK3 (clone 8G7^5^, in-house; Fig. 5a). These antibodies yielded specific signals via both immunofluorescence and immunoblot analyses (Fig. 5b-e). Interestingly, while phospho-RIPK3 concentrates into cytoplasmic clusters during necroptosis, this relocation event was not detected by other, anti-total RIPK3 antibodies (Fig. 5d). This finding suggests that the epitopes for RIPK3 detection are obscured via protein-protein interactions during the chronology of necroptotic signaling events. To further probe the steps in the pathway, we generated another mouse RIPK3 antibody, clone 1H12 (source: in-house), which has a favourable staining profile (Fig. 5b-c) and which recognises the translocation of mouse RIPK3 into cytoplasmic clusters during necroptosis (Fig. 5d). Notably, monoclonal antibodies raised against the N-terminal kinase domain of RIPK3, 8G7 and 1H12, detected several species of lower molecular weight than full length RIPK3 by immunoblot. These signals are likely to be spliced isoforms of mouse RIPK3 of varying lengths, each of which harbour the kinase domain antigen (residues 2-353), which are therefore not detected by antibodies directed towards the C-terminus of the full-length mouse RIPK3 isoform, such as ProSci 2283. In summary, there are several excellent options for the immunofluorescent detection of endogenous mouse RIPK3. Notably, unlike MLKL and RIPK1, the choice of fixative has no major bearing on the ability to detect mouse RIPK3 by immunofluorescence (Fig. 5c).

**Figure 5:**
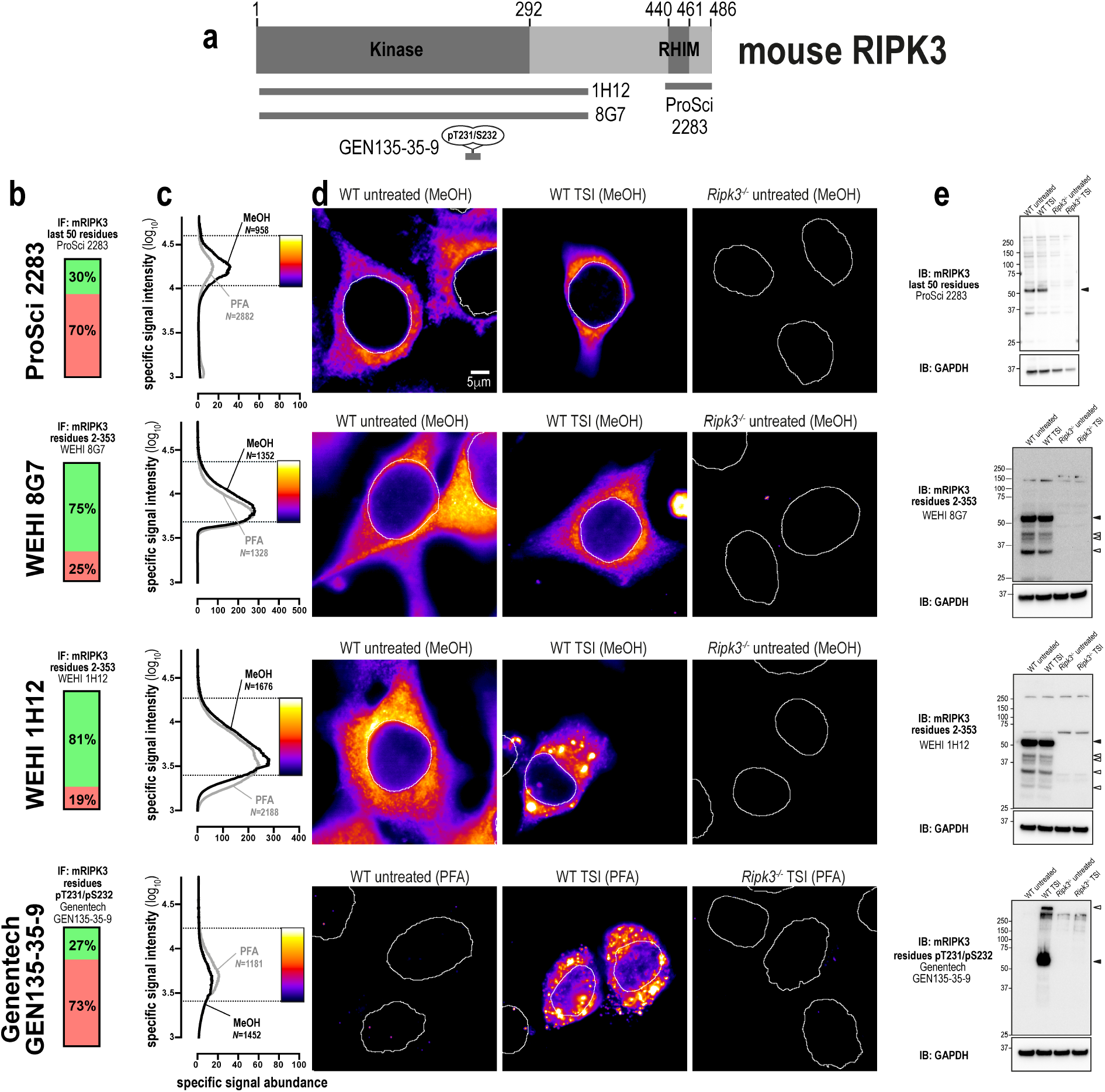
Unlike human RIPK3, mouse RIPK3 is highly amenable to detection by immunofluorescence and immunoblotting. **a** Mouse RIPK3 domain architecture showing the immunogens or epitopes for the tested anti-RIPK3 antibodies. **b** Quantitation of the percentage of gated signals for the tested RIPK3 antibodies. **c** Quantitation of specific signal abundance produced by the tested RIPK3 antibodies on methanol-fixed (MeOH) or paraformaldehyde-fixed (PFA) MDFs. The number of cells imaged (N) to generate each signal-to-noise curve is shown. **d** Micrographs of immunofluorescent signals for the tested RIPK3 antibodies on MDFs. As indicated by each pseudocolour look-up-table, only immunosignals within the respective gate in panel **c** were visualised. Data are representative of n=3 (in-house clone 8G7 and Genentech clone GEN135-35-9) and n=2 (ProSci 2283, in-house clone 1H12) independent experiments. Nuclei were detected by Hoechst 33342 staining and are demarked by white outlines in micrographs. **e** Immunoblot using the tested RIPK3 antibodies against lysates from wild-type and *Ripk3*^*-/-*^ MDFs. Closed arrowheads indicate the main specific band. Open arrowheads indicate other specific bands of interest. Immunoblots were re-probed for GAPDH as loading control.

### Immunofluorescent detection of mouse RIPK1

RIPK1 exhibits greater sequence identity between mouse and human orthologs than either RIPK3 or MLKL^41^. In keeping with this, both clone D94C12 (source: Cell Signaling Technology) and clone 38/RIP (source: BD Transduction Laboratories), which were raised against human RIPK1, also specifically detected mouse RIPK1 via immunofluorescence and immunoblotting (Fig. 6a-e). A polyclonal antibody raised against phospho-S166 of mouse RIPK1 (31122; source: Cell Signaling Technology) was also found to specifically detect RIPK1 via immunoblot (Fig. 6e) and in methanol-fixed, but not paraformaldehyde-fixed, cells undergoing necroptosis (Fig. 6b-d).

**Figure 6:**
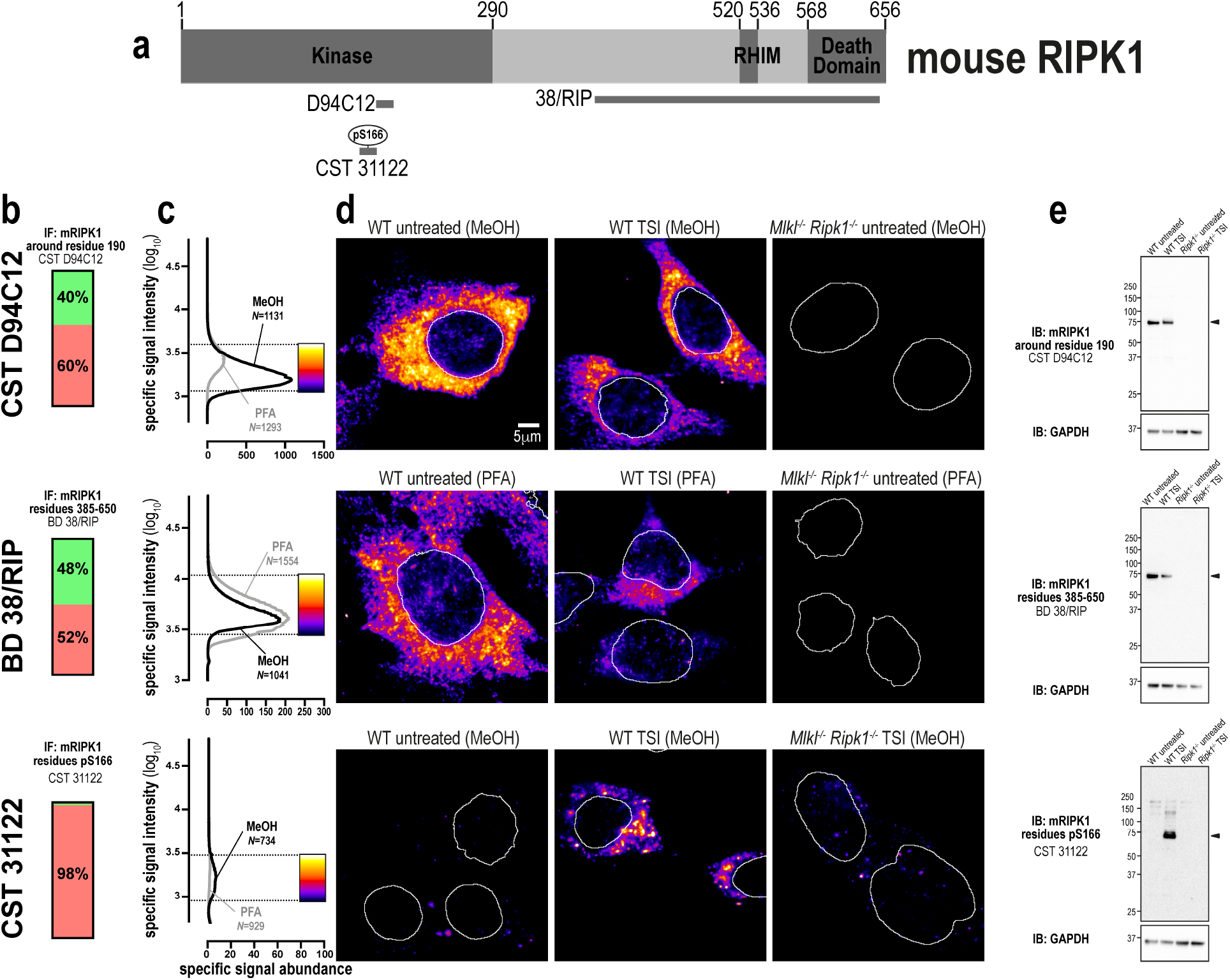
Three specific antibodies for imaging endogenous mouse RIPK1. **a** Mouse RIPK1 domain architecture showing the immunogens or epitopes for the tested anti-RIPK1 antibodies. **b** Quantitation of the percentage of gated signals for the tested RIPK1 antibodies. **c** Quantitation of specific signal abundance produced by the tested RIPK1 antibodies on methanol-fixed (MeOH) or paraformaldehyde-fixed (PFA) MDFs. The number of cells imaged (N) to generate each signal-to-noise curve is shown. **d** Micrographs of immunofluorescent signals for the tested RIPK1 antibodies on MDFs. As indicated by each pseudocolour look-up-table, only immunosignals within the respective gate in panel **c** were visualised. Data are representative of n=2 (Cell Signaling Technology clones D94C12 and 31122, and BD Transduction Laboratories clone 38/RIP) independent experiments. Nuclei were detected by Hoechst 33342 staining and are demarked by white outlines in micrographs. **e** Immunoblot using the tested RIPK1 antibodies against lysates from wild-type and *Ripk1*^*-/-*^ MDFs. Closed arrowheads indicate the main specific band. Open arrowheads indicate other specific bands of interest. Immunoblots were re-probed for GAPDH as loading control.

In summary, three highly-specific antibodies exist for mouse RIPK1. However, as was observed in human cells, necroptosis in mouse cells causes a marked reduction in RIPK1 levels (Fig. 6d-e). Thus, the specific immunosignals from the antibodies tested are of low intensity and abundance in necroptotic cells. Therefore, careful gating of signals is recommended for the study of mouse RIPK1 in necroptotic cells.

### Antibody cocktails for imaging endogenous necroptotic signaling

Having validated and compared the performance of numerous reagents, we now propose different antibody combinations that can be used to study endogenous necroptotic signaling (Fig. 7a-b). Because these recommendations are made to facilitate the investigation of endogenous necroptotic signaling in fixed cells, we also provide an ImageJ macro to generate signal-to-noise curves for the gating of specific immunosignals (Supplementary File 1). To exemplify the advantages of visualising endogenous necroptotic events *in situ*, we co-stained MDFs for non-phosphorylated MLKL (clone 5A6; source: in-house) and phosphorylated MLKL (clone D6E3G; source: Cell Signaling Technology). Similar to what was described for human MLKL^29^, mouse MLKL was observed to concentrate into cytoplasmic clusters during necroptosis. However, this compartmentalisation event is less pronounced than those observed for human MLKL in necroptotic HT29 cells (arrowhead; Fig. 7c). Also similar to what was described for human MLKL^29^, phosphorylated MLKL was observed to form focal structures at the plasma membrane, rather than docking uniformly with the cell periphery (Fig. 7c). Remarkably, although smaller than the hotspots of phosphorylated human MLKL that form in necroptotic HT29 cells, junctional accumulations of phosphorylated mouse MLKL were also apparent in necroptotic MDFs (Fig. 7d and Supplementary Video 1). These data suggest that the assembly of both mouse and human MLKL into macromolecular structures at the plasma membrane, and preferentially at sites of intercellular contact, may be an intrinsic and penultimate feature of necroptotic signaling.

**Figure 7:**
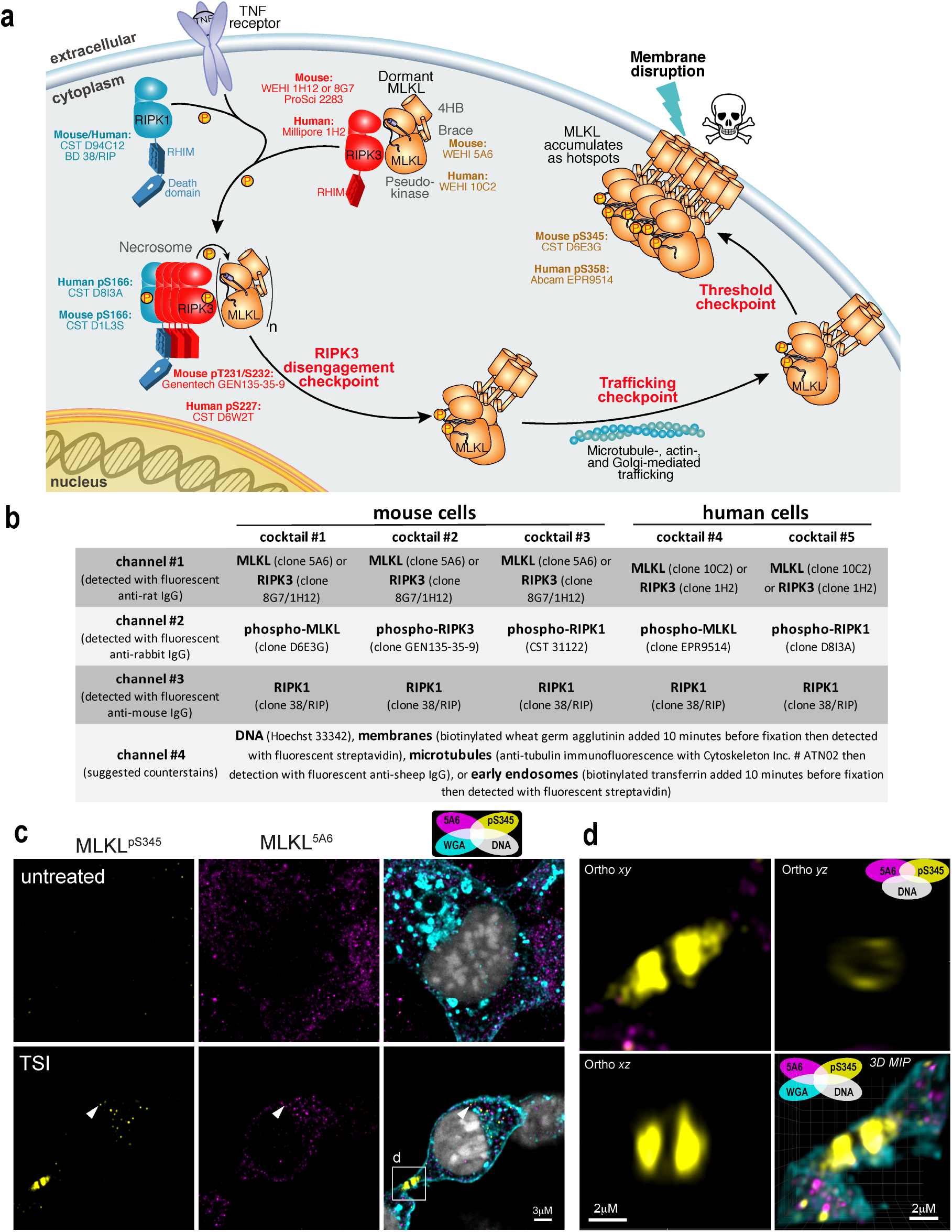
Optimised antibody cocktails for visualising endogenous necroptotic signaling in fixed human and mouse cells. **a** Cartoon summary of the currently-understood chronology of TNF-induced necroptosis. Recommendations of validated antibodies for immunostaining various steps in the necroptotic pathway are provided. **b** Summary of validated antibody cocktails and counterstains that can be multiplexed to examine endogenous necroptotic signaling in fixed human and mouse cells. Successful detection of specific signals relies on fixation in methanol, rather than crosslinking fixatives such as paraformaldehyde. **c** Two-dimensional Airyscan micrographs of Wheat Germ Agglutinin (WGA)-stained membranes, Hoechst-stained DNA and anti-MLKL immunosignals from clone 5A6 and clone D6E3G on methanol-fixed wild-type MDFs that had been left untreated or TSI-treated for 60min. Arrowhead exemplifies the small clusters of MLKL that form during necroptosis. The box indicates the junctional accumulation of phospho-MLKL. **d** Three-dimensional orthogonal projections and maximum intensity projection (MIP) of the boxed region from panel **c** showing a ring-like structure adopted by phospho-MLKL at the WGA-stained plasma membrane. Accompanied by Supplementary Video 1.

## DISCUSSION

Here, we profiled the immunofluorescent staining of MLKL, RIPK3 and RIPK1 in fixed human and mouse cells with a panel of 22 antibodies. While 17 of 22 antibodies were capable of detecting their respective necroptotic protein target in fixed wild-type cells and not their knockout counterparts, only 14 of 22 antibodies exhibited favourable signal-to-noise for robust immunostaining (as summarized in Fig. 7a).

The choice of fixative was a critical variable for the immunofluorescent detection of necroptotic proteins. With the exception of phospho-S227 in human RIPK3 and pT231/pS232 in mouse RIPK3, all other epitopes exhibited increased immunoreactivity in methanol-fixed samples. Indeed, immunodetection of MLKL was often found to be entirely contingent upon the use of methanol as a fixative. This observation has ramifications for the experimental and clinical immunohistological detection of cell death, which is almost exclusively performed on aldehyde-fixed samples. Thus, stringent antigen retrieval steps to reverse crosslinks will likely be necessary for the development of a diagnostic assay for necroptosis in patient samples. High resolution microscopy is also mostly performed on aldehyde-fixed cell monolayers. However, as these cell monolayers cannot typically withstand antigen retrieval, we recommend the use of methanol fixation for high resolution immunofluorescence of necroptosis; even though methanol fixation can be sub-optimal for both the retention of certain biomolecules^54^ and the preservation of cellular architecture^55, 56^. Indeed, all but one of the antibodies (source: Cell Signaling Technology clone D6W2T) in the suggested toolbox in Fig. 7a are compatible with methanol fixation.

Our data underscore the importance of using controlled experimental conditions to maximise the identification of specific immunosignals. To this end, we used wild-type cells that express high levels of MLKL, RIPK3 and RIPK1, and which have a well-characterised response to necroptotic stimuli. If possible, cell cultures should be fixed when low, but measurable, levels of necroptotic death are occurring (Fig. S1). This is because end-stage dead cells are often lost during/after methanol fixation, and because the immunosignals for phospho-RIPK1, phospho-RIPK3 and phospho-MLKL peak prior to cell death^29^. Notably, the use of knockout cells (for non-phospho-targets) or unstimulated wild-type cells (for phospho-sites) as a negative control is critical, as it provides the information needed to define the signal-to-noise profile of an antibody (Fig. 1b). Additionally, it is also helpful if different antibodies directed toward the same target yield concordant immunofluorescent signals, and if accompanying immunoblot data confirm their specificity. This combined approach was important for deducing that clone D6E3G (source: Cell Signaling Technology) is the best option for detecting phospho-MLKL in mouse cells (Fig. 4).

It is concerning that the specificity of some antibodies could not be verified. For instance, while numerous studies use the ProSci 2283 antibody to detect human RIPK3^57, 58, 59^, we found that it only recognised mouse RIPK3. Similarly, M6697 (source: Sigma-Aldrich) and clone 7C6.1 (source: Millipore) have been widely used to respectively detect human MLKL and mouse MLKL phospho-S345 ^39, 60, 61, 62, 63, 64, 65^, however these reagents exhibited a low degree of specificity in our hands. Nonetheless, as we did not exhaustively test the performance of these antibodies for immunofluorescence using a wide variety of fixatives, blocking agents, and detergents, we cannot categorically state that they are non-selective. Moreover, as ProSci 2283 (reported to detect mouse and human RIPK3) and Sigma-Aldrich M6697 (reported to detect human MLKL) are polyclonal antibodies, batch-to-batch variation may explain differences in their specificity between studies. It is also noteworthy that the majority of monoclonal antibodies that yielded the most abundant and specific signals were raised against recombinant full-length protein or component domains, rather than unfolded partial domains or peptides. We reasoned that epitopes that are exposed in the cellular forms of these proteins would be similarly presented within a folded protein for immune recognition in the host. Phosphosite-specific antibodies differ in this regard; these were raised using linear phosphopeptides and their specificity likely reflects the unstructured and solvent-exposed nature of the sequences in which they reside.

Broadly, the specific immunosignals described in this study provide consensus about three key compartmentalisation events that underlie necroptotic signaling^66^. First, the core necroptotic proteins - MLKL, RIPK3 and RIPK1 - predominantly reside in the cytosol under basal conditions. Interestingly, rather than being diffusely distributed across the cytosol, this necroptotic machinery exists as small puncta that are below the resolution limit of conventional light microscopy. It may be that these necroptotic proteins are preassembled into a membrane-less organelle under basal conditions. Second, RIPK1, RIPK3 and MLKL then coalesce into large cytoplasmic clusters. These clusters preferentially locate to the perinuclear space and are necrosome-related as they co-stain for the presence of phospho-RIPK1, -RIPK3 and -MLKL (Fig. 1-7 and ref.^29^). It is important to note that the formation of these necrosomal clusters in TSI-treated HT29 cells occurs subsequent to the phosphorylation of MLKL^29^. While these clusters likely represent a higher-order form of the necrosome, the mechanisms that govern cluster formation and limit their overall size are currently unknown. Third, as necroptotic signaling continues, phospho-MLKL is actively trafficked away from the clusters towards the plasma membrane^26^ where it accumulates into supramolecular assemblies called “hotspots”^29^. Recent findings suggest the release of phospho-MLKL from clusters arises from MLKL undergoing a conformational change that accompanies disengagement from necrosomal RIPK3^40^. Our data support such a disengagement step because neither RIPK1 nor RIPK3 were observed to accumulate at the plasma membrane during necroptosis in mouse or human cells (Fig. 2-3 and 5-6). This disengagement step, and formation of supramolecular assemblies by both mouse and human MLKL at the plasma membrane, are likely to be two additional checkpoints in necroptotic signaling.

## METHODS

### Materials

Primary antibodies and the dilutions used for immunoblotting/immunofluorescence were: rat anti-human MLKL (clone 10C2; produced in-house^29^; 1:2000/1:400), rat anti-human MLKL (clone 7G2; produced in-house^29^; 1:2000/1:400), rabbit anti-phospho-S358 human MLKL (Abcam ab187091; clone EPR9514; 1:2000/1:200), rat anti-mouse, human, rat, horse MLKL (clone 3H1; produced in-house^37^ and available as Millipore MABC604; 1:2000/ N/A), mouse anti-phospho-T357 human MLKL (Novus Biological MAB9187; clone 954702; 1:1000/1:200), rabbit anti-human MLKL (Sigma-Aldrich M6697; 1:1000/1:200), rabbit anti-RIPK3 (ProSci #2283; 1:1000/1:200), rabbit anti-human RIPK3 (Novus Biological NBP2-24588; 1:1000/1:200), mouse anti-human RIPK3 (Novus Biological; clone 780115; 1:1000/1:200), rat anti-human RIPK3 (clone 1H2; produced in-house^5^ and available as Millipore MABC1640; 1:2000/1:400), rabbit anti-phospho-S227 human RIPK3 (Cell Signaling Technology; clone D6W2T; 1:2000/1:200), rabbit anti-mouse or human RIPK1 (Cell Signaling Technology; clone D94C12; 1:2000/1:200), mouse anti-mouse or human RIPK1 (BD Biosciences; clone 38/RIP; 1:1000/1:100), rabbit anti-phospho-S166 human RIPK1 (Cell Signaling Technology; clone D8I3A; 1:2000/1:200), rat anti-mouse MLKL (clone 5A6; produced in-house), rabbit anti-phospho-S345 mouse MLKL (Abcam; clone EPR9515(2); 1:2000/1:200), mouse anti-phospho-S345 mouse MLKL (Millipore MABC158; clone 7C6.1; 1:2000/1:200), rabbit anti-phospho-S345 mouse MLKL (Cell Signaling Technology; clone D6E3G; 1:2000/1:200), rat anti-mouse RIPK3 (clone 8G7; produced in-house^5^ and available from Millipore as MABC1595; 1:2000/1:400), rat anti-mouse RIPK3 (clone 1H12; produced in-house; 1:2000/1:400), rabbit anti-phospho-T231/S232 mouse RIPK3 (Genentech; clone GEN135-35-9^53^; 1:2000/1:400), rabbit anti-phospho-S166 mouse RIPK1 (Cell Signaling Technology 31122; 1:2000/1:200), mouse anti-GAPDH (Millipore MAB374; 1:2000/N/A). Secondary antibodies for immunoblotting were: horseradish peroxidase (HRP)-conjugated goat anti-rat IgG (Southern Biotech 3010-05), HRP-conjugated goat anti-mouse IgG (Southern Biotech 1010-05), and HRP-conjugated goat anti-rabbit IgG (Southern Biotech 4010-05). All secondary antibodies for immunoblotting were used at a dilution of 1:10000. Secondary immunofluorescence detection reagents were: AlexaFluor647-conjugated donkey anti-rabbit IgG (ThermoFisher Scientific A31573), AlexaFluor568-conjugated donkey anti-rabbit IgG (ThermoFisher Scientific A10042), AlexaFluor568-conjugated donkey anti-mouse IgG (ThermoFisher Scientific A10037), AlexaFluor-594 donkey anti-rat IgG (ThermoFisher Scientific A-21209), AlexaFluor488-conjugated donkey anti-rat IgG (ThermoFisher Scientific A21208). All secondary antibodies for immunofluorescence were used at a dilution of 1:1000. Bond-Breaker TCEP solution (ThermoFisher Scientific 77720). *N*-Ethylmaleimide (Sigma-Aldrich E3876).

### 5A6 and 1H12 antibody production

Antibodies were generated at the Walter and Eliza Hall Institute Monoclonal Antibody Facility by immunizing Wistar rats with recombinant full-length mouse MLKL^37^ for clone 5A6 or recombinant mouse RIPK3 residues 2-353 for clone 1H12 and the previously-reported clone 8G7^5^ that was expressed and purified from Sf21 insect cells using the baculovirus expression system, before splenocytes were fused with SP2/O mouse myeloid cells and arising hybridoma lines cloned. The specificity of clone 5A6 for mouse MLKL was validated via immunoblot analyses as exemplified in Fig. 4e and Fig. S3. The specificity of clone 1H12 for mouse RIPK3 was validated via immunoblot analyses as exemplified in Fig. 5e.

### Cell lines

HT29 cells were provided by Mark Hampton (University of Otago). *RIPK1*^*-/-*^, *RIPK3*^*-/-*^ and *MLKL*^*-/-*^ HT29 cells have been previously reported^28, 29, 67^. MDF lines were generated in-house from the tails of wild-type C57BL/6J, *Mlkl*^*-/-*37^ and *Ripk3*^*-/-*68^ mice and immortalized by SV40 large T antigen as reported previously^35, 37^. The sex and precise age of these animals were not recorded, although our MDFs are routinely derived from tails from 8-week-old mice. *Mlkl*^*-/-*^*Ripk1*^*-/-*^ MDFs were derived from the belly/back dermis of *Mlkl*^*-/-*^*Ripk1*^*-/-*^ E19.5 mice^16^ and immortalized as described above. MDF lines were generated in accordance with protocols approved by the Walter and Eliza Hall Institute of Medical Research Animal Ethics Committee. Cell line identities were not further validated, although their morphologies and responses to necroptotic stimuli were consistent with their stated origins. Cell lines were routinely monitored to confirm they were mycoplasma-free.

### Cell culture

HT29 cells were maintained in Dulbecco’s Modified Eagle Medium (DMEM; Life Technologies) containing 8% v/v heat-inactivated fetal calf serum (FCS), 2 mM L-Glutamine/- GlutaMAX (ThermoFisher Scientific 35050061), 50 U ml^-1^ penicillin and 50 U ml^-1^ streptomycin (G/P/S) under humidified 10% CO2 at 37°C.

### Cell Treatment

Cells were seeded into Ibi-treated 8-well μ-slides (Ibidi 80826) in media containing 8% v/v FCS and G/P/S at 3.0 × 10^4^ cells per well for HT29 and 0.25 x 10^4^ cells per well for MDFs. Cells were left to adhere overnight then treated in media containing 1% v/v FCS and G/P/S and supplemented with the following stimuli: 100ng/mL recombinant human TNF-*α*-Fc (produced in-house as in ref.^69^), 500nM Smac mimetic/Compound A (provided by Tetralogic Pharmaceuticals; as in ref.^70^), 5μM IDN-6556 (provided by Tetralogic Pharmaceuticals). Unless stipulated, HT29 cells were stimulated for 7.5 h with TSI and MDF cells were stimulated for 90 min with TSI.

### LDH release

Colorimetric LDH release assay kit (Promega G1780) was performed according to manufacturer’s instructions.

### Whole cell lysate, SDS-PAGE, immunoblot & quantification

Cells were lysed in ice-cold RIPA buffer (10mM Tris-HCl pH 8.0, 1mM EGTA, 2mM MgCl_2_, 0.5% v/v Triton X100, 0.1% w/v Na deoxycholate, 0.5% w/v SDS and 90mM NaCl) supplemented with 1x Protease & Phosphatase Inhibitor Cocktail (Cell Signaling Technology 5872S) and 100U/mL Benzonase (Sigma-Aldrich E1014). Whole cell lysates were boiled for 10 minutes in 1× SDS Laemmli sample buffer (126 mM Tris-HCl, pH 8, 20% v/v glycerol, 4% w/v SDS, 0.02% w/v bromophenol blue, 5% v/v 2-mercaptoethanol), and resolved on 1.5mm NuPAGE 4–12% Bis-Tris gels (ThermoFisher Scientific NP0335BOX) using MES Running buffer (ThermoFisher Scientific NP000202) or Bio-Rad Criterion TGX 4-15% gels (Bio-Rad 5678085) using 1x TGS buffer (Bio-Rad 1610772). After transfer onto nitrocellulose, membranes were blocked in 5% w/v skim milk powder in TBS-T, probed with primary antibodies (see ***Materials*** above) then the appropriate HRP-conjugated secondary antibody (see ***Materials*** above) and signals revealed by enhanced chemiluminescence (Merck P90720) on a ChemiDoc Touch Imaging System (Bio-Rad). Between each probe, membranes were incubated in stripping buffer (200mM glycine pH 2.9, 1% w/v SDS, 0.5mM TCEP) for 30 minutes at room temperature then re-blocked.

### Subcellular fractionation, BN-PAGE & immunoblot

HT29 cells or MDFs were seeded into 6-well plates (1.0 × 10^6^ cells per well) in media containing 8% v/v FCS and G/P/S and equilibrated overnight under humidified 10% CO2 at 37 ^°^C conditions. Cells were then treated in media containing 1% FCS and G/P/S supplemented with the agonists (as indicated above). Cells were fractionated into cytoplasmic and membrane fractions^35^. Cells were permeabilized in MELB buffer (20 mM HEPES pH 7.5, 100 mM KCl, 2.5 mM MgCl_2_ and 100 mM sucrose, 0.025% v/v digitonin, 2 μM N-ethyl maleimide, phosphatase and protease inhibitors). Crude membrane and cytoplasmic fractions were separated by centrifugation (5 minutes 11,000*g*), and fractions prepared in buffers to a final concentration of 1% w/v digitonin. The samples were resolved on a 4–16% Bis-Tris Native PAGE gel (ThermoFisher), transferred to polyvinylidene difluoride (Merck IPVH00010). After transfer, membranes were destained (in 50% (v/v) methanol, 25% (v/v) acetic acid), denatured (in 6M Guanidine hydrochloride, 10 mM Tris pH6.8, 5 mM *β*-mercaptoethanol), blocked in 5% skim milk (Diploma), and probed in the same manner as above.

### Protein production and purification

Full-length human and mouse MLKL and the C-terminal, pseudokinase domain of mouse and human MLKL were expressed in Sf21 insect cells using the Bac-to-Bac system (ThermoFisher Scientific) and purified using established procedures^28, 37, 42^. Briefly, proteins were expressed with an N-terminal, TEV protease-cleavable GST (full length human MLKL) or His6 (full length mouse MLKL and pseudokinase domains) tags and captured from lysates using glutathione resin (UBP Bio) or HisTag Ni-NTA resin (Roche) respectively. Proteins were cleaved on-resin from the GST tag or off-resin for His6 tags using His-tagged TEV protease, before protease was removed by Ni-NTA chromatography (HisTag resin, Roche). Protein was concentrated by centrifugal ultrafiltration and applied to a Superdex-200 (GE Healthcare) size exclusion chromatography column; protein was eluted in 20 mM HEPES pH 7.5, 200 mM NaCl, 5% v/v glycerol. Protein containing fractions were spin concentrated to 5-10 mg/mL, aliquoted, snap frozen in liquid N2 and stored at –80°C until required.

### Immunofluorescence

Cells in 8-well μ-Slides (Ibidi 80826) were ice-chilled for 3 minutes, then washed in ice-cold Dulbecco’s PBS (dPBS; ThermoFisher Scientific 14190144), then fixed for 30 minutes in either ice-cold methanol or ice-cold 4% w/v PFA. Cells were washed twice in ice-cold dPBS, then blocked in ice-cold Tris-balanced salt solution with 0.05% v/v Triton-X100 (TBS-T) supplemented with 10% v/v donkey serum (Sigma-Aldrich D9663) for >1 hour. Cells were incubated in primary antibodies (see ***Materials*** above) overnight at 4 °C in TBS-T with 10% v/v donkey serum. Cells were washed twice in TBS-T then incubated in the appropriate secondary antibodies supplemented with 0.1μg/mL Hoechst 33342 (ThermoFisher Scientific H3570) for 3 hours at room temperature with gentle rocking. Cells were washed four times in ice-cold TBS-T then stored at 4 °C until imaged. Where indicated, to demarcate the plasma membrane, 2μL of biotinylated wheat germ agglutinin (Sigma-Aldrich L5142) was added to each well 10 minutes before fixation. The fixed wheat germ agglutinin was then detected via the addition of 1:1000 dilution of DyLight650-conjugated streptavidin (ThermoFisher Scientific 84547) during the secondary antibody incubation step.

### Two-dimensional epifluorescence microscopy

Samples in TBS-T were imaged on an Inverted Axio Observer.Z1 microscope (Zeiss) with the following specifications: C-Apochromat 40x/1.20 W autocorr UV VIS IR lens, HXP 120V excitation source, AlexaFluor647 and DyLight650 imaged with a λ_Excitation_=625-655nm; λ_beamsplitter_=660nm; λ_Emission_=665-715nm filter, AlexaFluor568 imaged with a λ_Excitation_=532-544nm; λ_beamsplitter_=560nm; λ_Emission_=573-613nm, AlexaFluor488 imaged with a λ_Excitation_=450-490nm; λ_beamsplitter_=495nm; λ_Emission_=500-550nm, Hoechst 33342 imaged with a λ_Excitation_=359-371nm; λ_beamsplitter_=395nm; λ_Emission_=397-∞ nm, a sCMOS PCO.edge 4.2 camera, ZEN blue 2.5 pro capture software and ImageJ 1.53c post-acquisition processing software^71^. Typically, for each independent experiment, 5-10 randomly selected fields were captured per treatment group, whereby only the Hoechst 33342 signal was visualised prior to multi-channel acquisition. To ensure consistent signal intensities across independent experiments, the same excitation, emission and camera settings were used throughout this study.

### Airyscan microscopy

Fixed immunostained cells in 8-well μ-Slides (Ibidi 80827) were subjected to super-resolution 3-dimensional Airyscan microscopy on an Inverted LSM 880 platform (Zeiss) equipped with the following specifications: a 63x/1.4 N.A. PlanApo DIC M27 oil immersion objective (Zeiss), 405-, 488-, 568- and 640-nm laser lines, and radially-stacked Airyscan GaASP detectors set to SR-mode, 405-, 488-, 561-, 633-nm laser lines and ZEN black 2.3 SP1 FP3 v14.0 capture software. Image stacks were acquired with a z-step size of 159nm. Super-resolution deconvolution was performed using the automated ‘3D AiryScan Processing’ function of ZEN blue software.

### Hotspot quantitation

Micrographs (captured as in ***two-dimensional epifluorescence microscopy***) were opened in ImageJ 1.53c^71^. A rolling ball filter of 7 was applied and phospho-S358 MLKL immunosignals thresholded (*≥*7000 units) and objects segmented using the ‘Analyze>Analyze Particles’ tool. Segmented objects with size 0.5-100μm^2^ and feret diameter>2 (i.e. elliptical objects) were considered hotspots. The number of segmented objects per 100 cells was taken as an index of hotspot occurrence. The mean size of the segmented objects was taken as an index of hotspot size. The fluorescence intensity of each segmented object divided by its size was taken as an index of hotspot intensity.

## ACKNOWLEDGEMENTS

We thank the Walter and Eliza Hall Institute Monoclonal Antibody Facility for their assistance generating clones 3H1, 10C2, 7G2, 1H2, 5A6, 8G7 and 1H12; and Cathrine Hall for assistance generating *Mlkl*^*-/-*^ *Ripk1*^*-/-*^ MDF lines. We are grateful to the Australian National Health and Medical Research Council for fellowship (J.M.H., 1142669; E.D.H., 1159488; J.M.M., 1172929), grant (1124735, 1124737, and 1105023), and infrastructure (IRIISS 9000587) support, with additional support from the CASS Foundation (A.L.S.) and the Victorian Government Operational Infrastructure Support scheme. We acknowledge the support for A.V.J. from an Australian Research Training Program Scholarship.

## AUTHOR CONTRIBUTIONS

Investigation: A.L.S., K.P., C.F., J.M.H., L.W., J.R., A.V.J., C.R.H., X.G., S.N.Y. Supervision and Methodology: A.L.S., J.M.H., K.L.R., E.D.H. and J.M.M.; Conceptualization: A.L.S., E.D.H. and J.M.M.; Writing – Original Draft: A.L.S. and J.M.M.; Writing – Review & Editing: all authors.

## MATERIALS REQUESTS

All materials will be provided under Materials Transfer Agreement upon request to James Murphy.

