## Supplementary Information for "A toolbox for imaging RIPK1, RIPK3 and MLKL in mouse and human cells"

Supplementary Figure 1

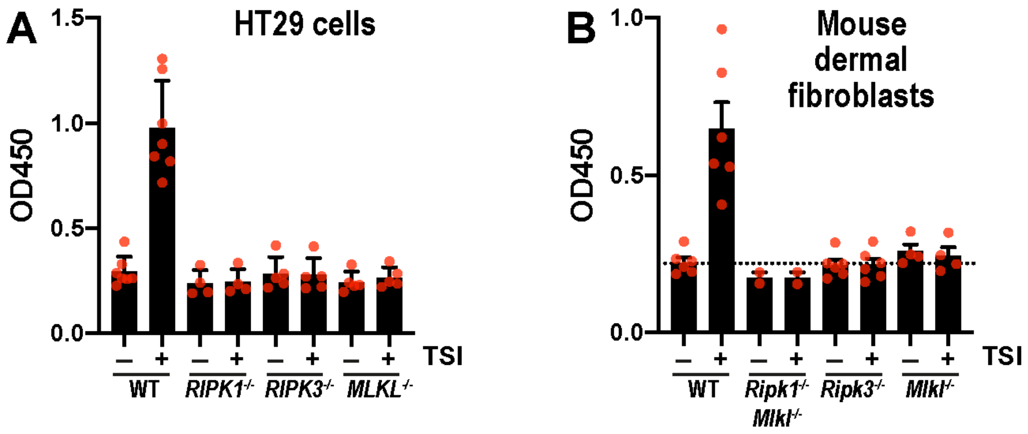


**Supplementary Figure 1. Necroptotic cell death was confirmed in samples before immunofluorescent staining.**

The release of lactate dehydrogenase into the conditioned media of HT29 cells (**A**) and MDFs (**B**) was measured prior to immunofluorescent staining as a measure of necroptotic cell death. Graphs plot the raw absorbance units from these lactate dehydrogenase release assays. Each red dot indicates the mean of one independent experiment and bars indicating the mean + S.E.M. across n=2-6 independent experiments. Only once TSI-induced necroptotic responses were verified in wild-type cells, but not in MLKL-, RIPK3- and RIPK1-deficient cells, these same samples were subjected to immunofluorescence.

### Supplementary Figure 2

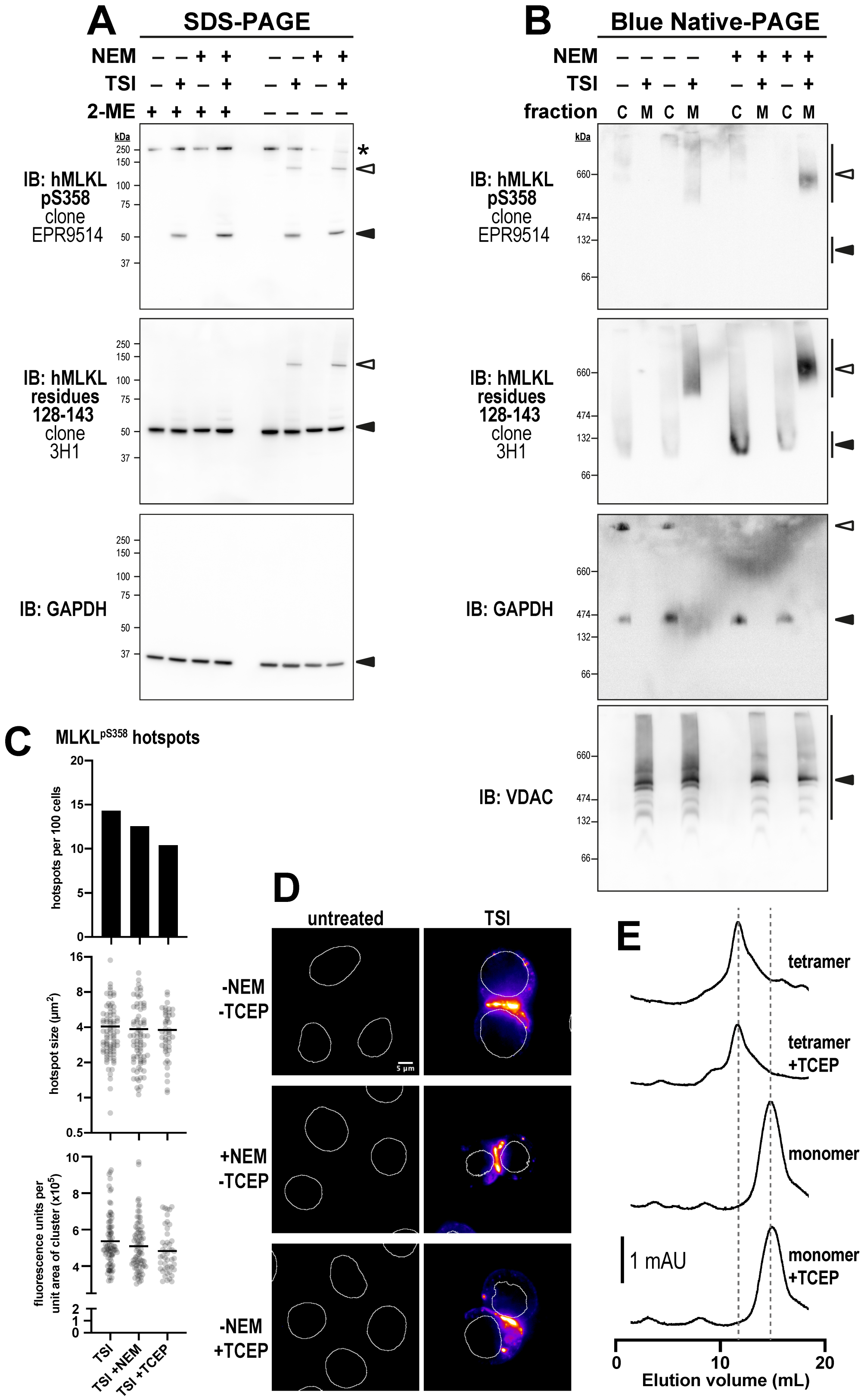

**Supplementary Figure 2. Only a fraction of oligomeric MLKL is disulfide-crosslinked prior to necroptotic cell death.**

**A** HT29 cells were treated for 4.5 hours, whole cell lysates were prepared in the absence/presence of 30 mM *N*-ethylmaleimide (NEM) then subjected to SDS-PAGE in the absence/presence 2-mercaptoethanol (2-ME) and immunoblotted for MLKL (clone 3H1), phospho-MLKL (clone EPR9514) and a GAPDH loading control. Closed arrowheads indicate the main specific bands. Open arrowheads indicate MLKL oligomers. Asterisks indicate non-specific bands that could otherwise confound data interpretation. Data are representative of n = 2 independent experiments. **B** The same experiment as in panel **a** except that cytosol (C) and membrane (M) fractions were prepared in the absence/presence of NEM then resolved by Blue Native-PAGE and immunoblotted for MLKL (clone 3H1; available as Millipore MABC604) and phospho-MLKL (Abcam clone EPR9514). Fractionation was verified by probing for voltage-dependent anion channel 1 (VDAC; membrane), GAPDH (cytosol). Data are representative of n = 2 independent experiments. **C-D** HT29 cells were TSI-treated for 7.5 hours, methanol-fixed in the presence/absence of 10μM NEM, treated in the presence/absence of 0.5mM Tris(2-carboxyethyl)phosphine (TCEP), immunostained for phospho-MLKL (clone EPR9514), DNA counterstained with Hoechst 33342 and then imaged via epifluorescence. Panel **C** shows the frequency, intensity and size of individual phospho-MLKL hotspots at the plasma membrane from one experiment (N=566 cells for TSI, N=613 cells for TSI+NEM and N=443 cells for TSI+TCEP). Population means are indicated by lines. Data are representative of n=2 independent experiments. Panel **D** shows representative micrographs of phospho-MLKL hotspots at the plasma membrane. Data are representative of n=2 independent experiments. **E** Recombinant wild-type human MLKL monomer or tetramer was incubated in the presence or absence of 1 mM TCEP and approximately 0.1 mg of protein was resolved by size exclusion chromatography using a Superdex

200 10/300 Increase column at a flow rate of 0.4 mL/min. Elution profiles are representative of n=2 independent experiments.

### Supplementary Figure 3

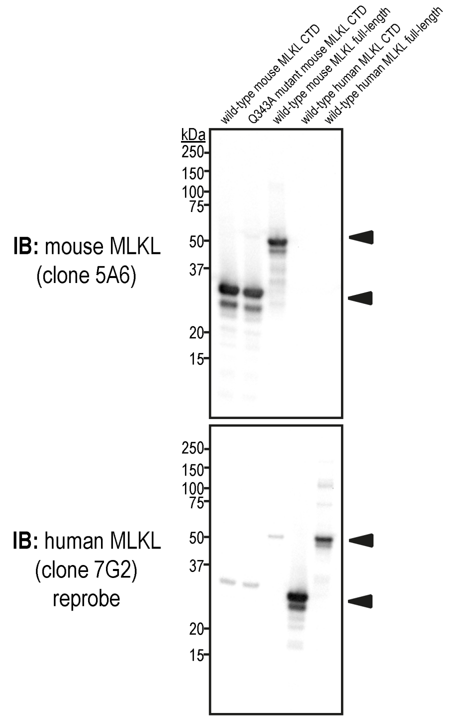


**Supplementary Figure 3. Clone 5A6 is a new monoclonal antibody that specifically recognises the C-terminal, pseudokinase domain of mouse MLKL.** 0.25 μg of each of the indicated recombinant proteins were fractioned by SDS-PAGE and immunoblotted with the anti-mouse MLKL antibody (5A6 clone), verifying that 5A6 specifically recognises the C-terminal pseudokinase domain (CTD) of mouse MLKL. Immunoblot was reprobed with human MLKL C-terminal pseudokinase domain-specific antibody, clone 7G2 (lower panel).

**Supplementary File 1 – Image J macro to facilitate the generation of signal-to-noise curves using immunofluorescence data.**

**Supplementary Video 1 – Mouse MLKL also accumulates at intercellular junctions.** Three-dimensional maximum intensity projection of a ring-like junctional structure adopted by phospho-MLKL at the plasma membrane. Wheat germ agglutinin-stained membrane in cyan, $MLKL^{5A6}$ immunosignal in magenta and $MLKL^{pS345}$ immunosignal in yellow.
